# Coarse-grained simulations of actomyosin rings point to a nodeless model involving both unipolar and bipolar myosins

**DOI:** 10.1101/194910

**Authors:** Lam T. Nguyen, Matthew T. Swulius, Samya Aich, Mithilesh Mishra, Grant J. Jensen

## Abstract

Cytokinesis in most eukaryotic cells is orchestrated by a contractile actomyosin ring. While many of the proteins involved are known, the mechanism of constriction remains unclear. Informed by existing literature and new 3D molecular details from electron cryotomography, here we develop 3D coarse-grained models of actin filaments, unipolar and bipolar myosins, actin crosslinkers, and membranes and simulate their nteractions. Exploring a matrix of possible actomyosin configurations suggested that node-based architectures ike those presently described for ring assembly result in membrane puckers not seen in EM images of real cells. Instead, the model that best matches data from fluorescence microscopy, electron cryotomography, and biochemical experiments is one in which actin filaments transmit force to the membrane through evenly-distributed, membrane-attached, unipolar myosins, with bipolar myosins in the ring driving contraction. While at this point this model is only favored (not proven), the work highlights the power of coarse-grained biophysical simulations to compare complex mechanistic hypotheses.

**Significance Statement:** In most eukaryotes, a ring of actin and myosin drives cell division, but how the elements of the ring are arranged and constrict remain unclear. Here we use 3D coarse-grained simulations to explore various possibilities. Our simulations suggest that if actomyosin is arranged in nodes (as suggested by a popular model of ring assembly), the membrane distorts in ways not seen experimentally. Instead, actin and myosin are more ikely uniformly distributed around the ring. In the model that best fits experimental data, ring tension is generated by interactions between bipolar myosins and actin, and transmitted to the membrane via unipolar myosins. Technologically the study highlights how coarse-grained simulations can test specific mechanistic hypotheses by comparing their predicted outcomes to experimental results.

## Introduction

It is well known that an actomyosin ring (AMR) drives cell division in most eukaryotic cells, but how it contracts and how force is transmitted to the membrane remain unclear (1, 2). Two components involved in contraction are actin filaments (F-actin) and the motor protein, non-muscle myosin II, which exerts tensile force on F-actin through a processive ATP-dependent power stroke mechanism (3). Both proteins are essential for cytokinesis and localize to an equatorial contractile ring during mitosis (4–12). Fluorescence studies of ring assembly in *Schizosaccharomyces pombe*, a rod-shaped unicellular fission yeast that shares most of the cytokinesis genes with metazoans (1), showed that the ring components first form a broad band of nodes (13, 14) that coalesce into a ring at the division plane (15). Recent electron cryotomography (ECT) of dividing fission yeast showed, however, that F-actin termini are apparently randomly distributed around the ring (16), calling into question whether nodes continue to exist during constriction. F-actin in the contractile ring is contributed by both existing actin cables (17) and de novo nucleation, primarily by the formin Cdc12p (18), a barbed-end actin-capping dimeric protein that is essential for ring assembly in fission yeast (19). While it has been proposed that ring tension is transmitted to the membrane via connection between the actin barbed end and Cdc12p, which either exists individually (2) or at nodes (20), this mechanism has not been proven.

There are two myosin type-II heavy chains (Myo2p and Myp2p) in the contractile ring. Myo2p, the essential type II myosin (6, 21), plays the leading role in ring assembly while the second, non-essential, unconventional type II myosin, Myp2p, is the major driver for ring constriction (22), consistent with its arrival at the division site immediately prior to ring constriction (12). Recent evidence indicates that during constriction, Myo2p and Myp2p are distributed in two distinct concentric rings (22), but the causes and functional implications of this segregation are unknown. While previous simulation studies have described myosin as bipolar (23, 24), and this assumption is supported by some *in vitro* evidence (25, 26), myosin has also been proposed to exist in a unipolar form with its C-terminal tail tethered to the membrane and its N-terminal motor domain in the cytoplasm, in a ‘bouquet-like’ arrangement (20, 27). Further study is needed to elucidate how myosin is organized within the ring and how it generates tension during constriction.

In addition to F-actin and myosin, the actin crosslinkers *α*-actinin and fimbrin have been reported to be important for assembly of the ring (28, 29). While α-actinin is present in the ring during constriction, it is not clear whether fimbrin is present as well (30). In vitro, however, addition of actin-crosslinkers stalls ring contraction (31). Thus it is presently unclear how these actin crosslinkers affect ring contraction. Cofilin has also been reported to help maintain the structure of the ring, but its seemingly counterintuitive function as an F-actin severing protein (32, 33) leaves its role during ring constriction unclear.

Simulations have been used previously to explore constriction of the actomyosin ring (34). In an early continuum model, discrete molecules were not described. Instead the ring was represented by density values and the roles of myosin and crosslinkers were implicitly represented using coefficients of tension contribution (35). Simulations based on this model suggested that actin depolymerization in the presence of end-tracking crosslinkers could drive constriction, but whether such a crosslinker exists is unknown. Later simulations further explored this same idea, modeling individual filaments as lines with defined polarity (36). In more recent work, the ring was modeled as a 2D band in which actin filaments were modeled as chains of beads and clusters of myosins were represented as single beads which exerted force on actin filaments in close proximity (24). Parameters were found in which this 2D model produced tension similar to that measured in fission yeast protoplasts. Simulations have also explored the condensation of the ring before constriction (37).

Prompted by new electron cryotomography (ECT) data revealing for the first time the native 3D organization of the actin filaments and the membrane in dividing yeast cells (16), here we developed more detailed and 3D coarse-grained simulations to explore different hypotheses about how actin and myosin might constrict the membrane. F-actin, unipolar and bipolar myosins, and actin crosslinkers were all modeled using a bead-spring representation. A flexible cylindrical membrane was also modeled. To make actomyosin interactions as realistic as possible, the ATPase cycle of myosin was implemented in step-by-step detail. Random forces were further added to mimic thermal fluctuation.

First, we introduced the basic components of the ring one-by-one to define a minimal set of components and rules necessary for constriction. In doing so, we found that actin crosslinkers are required to propagate tension through the ring, and that introducing cofilin to sever bent F-actin helps reproduce the filament straightness observed by ECT. We then explored sixteen candidate actomyosin architectures and ring-to-membrane attachments. Combined with ECT data, our results suggest that actomyosin does not exist in nodes during constriction. Judged by all currently available experimental data, our simulations favor a model in which the ring tension is generated primarily through interactions between bipolar myosins and actin filaments, and is transmitted to the membrane via unipolar myosins, which are individually attached to the membrane. Due to the 3D and dynamic nature of our data, which is much better presented in movies than static figures, we encourage readers to begin by watching Movie S1, which presents (i) the elements and properties of our 3D coarse-grained model of the contractile ring, (ii) building the initial model, (iii) exploration of different actomyosin configurations, and (iv) a final model that best agreed with experimental data.

## Results

### Basic components of the ring

To build a coarse-grained model of the contractile ring, three main components of the ring including F-actin, myosin and crosslinkers were represented using a bead-spring model (Fig. 1A). Each filament was modeled as a chain of beads connected by springs, each myosin was modeled to be either unipolar or biopolar, and each crosslinker was modeled to have two actin binding domains at the two ends. The membrane was modeled as a sheet of beads, originally having a cylindrical shape (Fig. 1A). Actin-myosin interaction was modeled to occur in a power-stroke fashion in which the myosin ATPase cycle had five steps (Fig. 1B). The power stroke was generated via changing the angle of the myosin head as it transitioned between its ATPase phases (see **Methods** for details).

**Figure 1:**
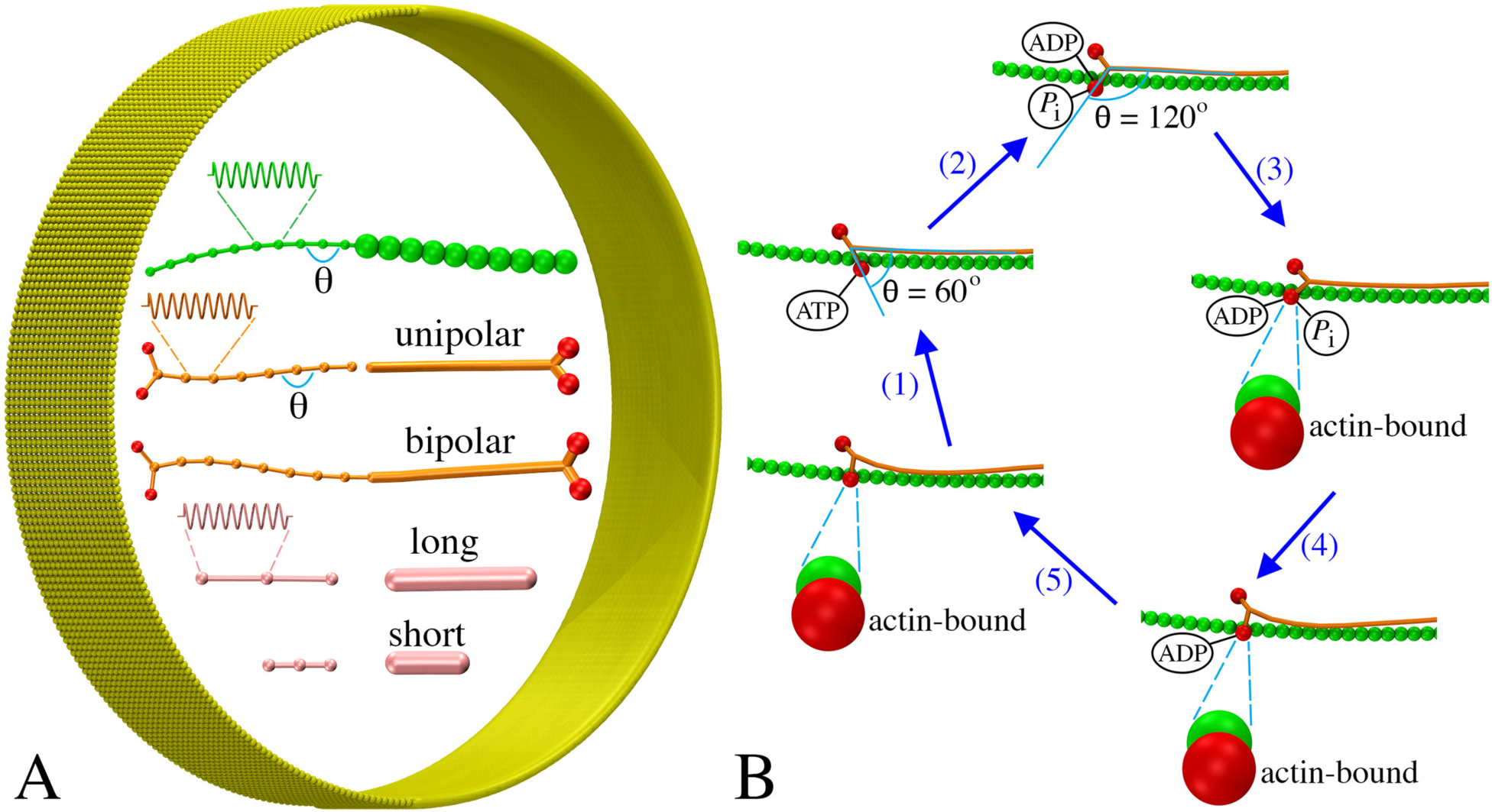
Coarse-graining the actomyosin system: (A) Models of F-actin (green), myosins (tail in orange, heads in red), actin crosslinkers (pink), and membrane (yellow) (see text for details). Note the same visualizations and colors in the right are used for all following figures unless otherwise stated. (B) The ATPase cycle of myosin was modeled in five steps: myosin (1) binds ATP and releases actin, (2) hydrolyzes ATP, (3) binds actin, (4) releases phosphate, and (5) releases ADP.

Many proteins are present at the mid-cell during constriction, but it is unclear which are essential for the contractility of the ring. We therefore started with a very simple model, testing interactions between bipolar myosin and F-actin of mixed polarities, originally arranged into a ring (**Methods/Initial ring configuration**). In this test, a membrane was added to confine the actomyosin system, but membrane constriction was not expected since it was not linked to the ring (Fig. 2A). As myosin moved along F-actin toward their plus ends in a ATPase-dependent power-stroke fashion (Fig. 1B), the filaments slid, bent and oriented randomly, but the ring did not constrict due to the lack of long-range propagation of tension around the ring (Fig. 2B; Movie S1, at 2:10). Reasoning that crosslinking F-actin would help propagate tension, actin crosslinkers were added, and the ring began to contract, despite losing the original ring-like arrangement of F-actin (Fig. 2C; Movie S1, at 2:35). Linking the ring to the membrane (**Methods/Membrane tethering**) resulted in membrane constriction showing that a ring composed of F-actin, myosin and actin crosslinkers is capable of generating tension and constricting the membrane (Fig. 2D; Movie S1, at 3:01). As the membrane was pulled inward, cell wall material was added behind preventing the membrane from relaxing back (see **Methods**). The ring-like arrangement of F-actin was now maintained, suggesting that membrane attachment contributes to maintenance of the ring structure. Note that in later simulations of model 1, tethering the actin plus end and unipolar myosin tail to membrane-bound nodes produced tension temporarily in the absence of crosslinkers. As the nodes were able to slide on the membrane to aggregate into separated large clusters, however, the ring was quickly broken (SI Appendix/Fig. S2A), pointing again to the need of crosslinkers for ring constriction.

**Figure 2:**
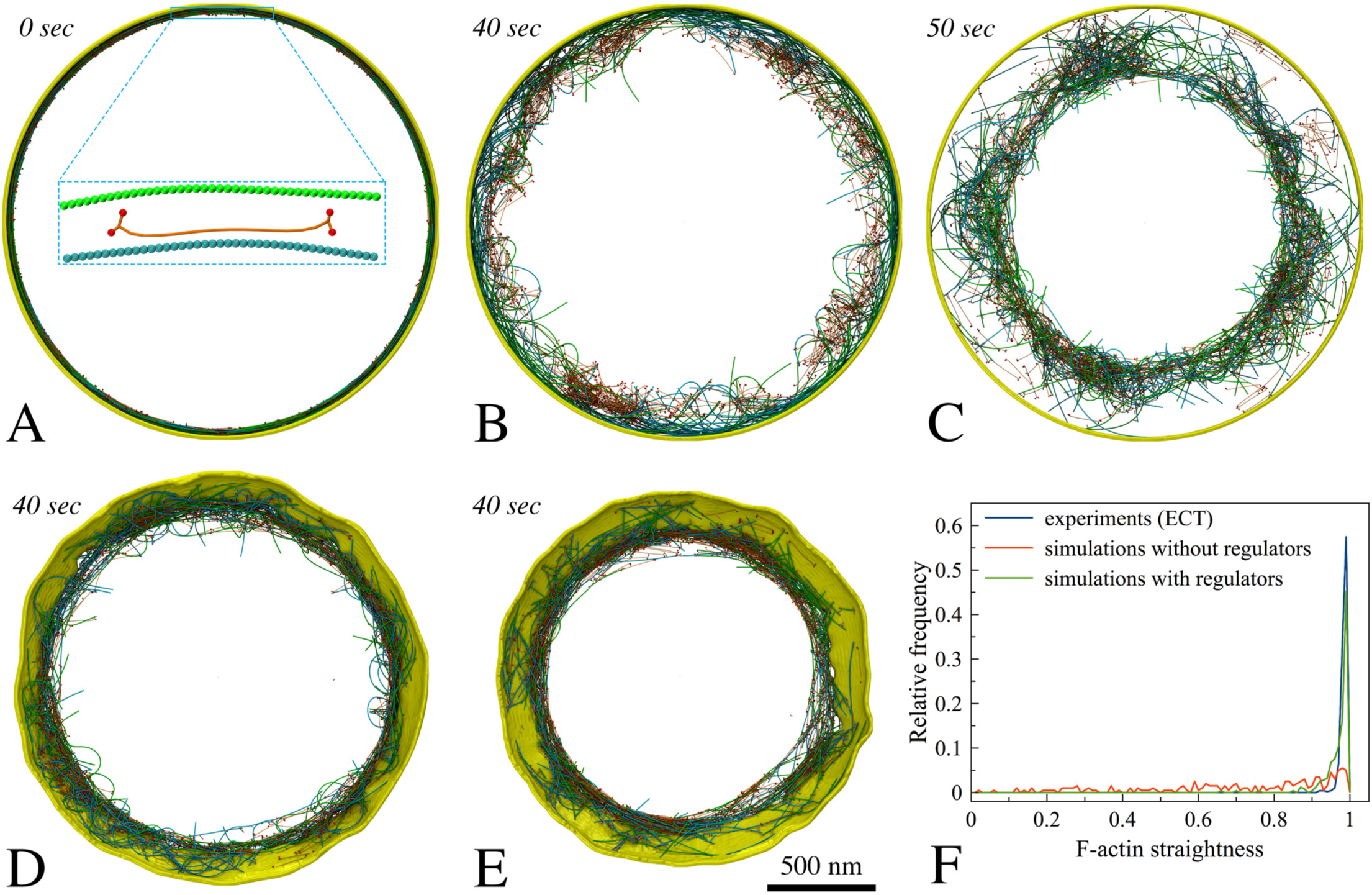
Setting up basic components of the simulated constriction system: F-actin (opposing polarities in green and cyan), bipolar myosin, crosslinkers, and membrane. Italic fonts indicate simulation times. (A) The initial ring was composed of F-actin, bipolar myosin and membrane. (B) The actomyosin ring did not contract in the absence of crosslinkers. (C) In the presence of crosslinkers, the ring did contract. (D) Adding tethers between F-actin and the membrane caused the ring to constrict the membrane. (E) Adding regulatory factors (tension-dependent interaction between actin and myosin, filament orientation-dependent crosslinking, cofilin function, and actin turnover) reduced F-actin bending. (F) Histogram of straightness factors for filaments visualized by ECT (16) (blue), simulations without regulatory factors (red), and with regulatory factors (green).

### F-actin straightness regulatory factors

At this stage, the simulated F-actin did not mimic the consistently straight filaments observed experimentally (16), but they were highly bent (Fig. 2D, F, SI Appendix/Fig. S1). To study how the myosin processivity would influence bending, we reduced the myosin duty ratio (see **Methods/Myosin ATPase cycle** for the definition). As the first step of the ATPase cycle was slowed down 5 and 10 times, the duty ratio was reduced from the original value of ~ 0.72 to 0.35 and 0.21, slowing down ring constriction and delaying filament bending, but this did not eliminate bending. Inspecting the simulation results, we identified at least two factors that contributed to filament bending. First, if an F-actin was crosslinked close to its minus end while myosin was walking toward its free plus end, the plus end was pulled toward the minus end, bending the filament (Movie S1, at 3:42). As one proposed ability of F-actin is tension sensing (38), and myosin is known to bind preferentially to F-actin under tension (39), we added a rule that myosin could bind to actin only if the filament was crosslinked upstream. Note that even if we had tracked them in the simulation, other binding events would not have contributed tension since loose filaments simply move when pulled.

Second, if an actin filament had each of its ends crosslinked to two different filaments sliding toward one another, the filament would bend (Movie S1, at 4:03). We reasoned that bending was not seen *in vivo* because either (i) crosslinks were released on the bent filament or (ii) the filament was broken. Hypothesizing that torque facilitates crosslink release, we added a rule that the probability of crosslink release increases with the angle between two filaments at their crosslinked location (**Methods/Torque-facilitated crosslinker release** for details). Next, considering that the actin-depolymerizing factor cofilin preferentially severs F-actin that is not under tension (40), we introduced its function into the simulation by stipulating that the probability of filament breaking increases with bending angle (**Methods/Cofilin function** for details).

Another factor that might affect F-actin bending is actin depolymerization, which has been shown to occur rapidly during constriction (41). Actin turnover was therefore added (**Methods/Protein turnover** for details). Further, turnover of myosin and crosslinkers was also implemented (**Methods/Protein turnover** for details) since this occurs in fission yeast (29, 41, 42). In the presence of these regulatory rules, F-actin bending was prevented *in silico* (Movie S1, at 5:04; Fig. 2E, F), thus recapitulating the filament straightness observed experimentally (16).

### Exploration of actomyosin architecture models

Having established a working core model, we explored fifteen plausible configurations and arrangements of F-actin and myosin to study how they would constrict the membrane (Fig. 3). We reasoned that the membrane must be tethered to either actin or myosin, or both, to enable membrane constriction. The four configurations of actin are illustrated in Fig. 3 (panels A1–A4). In (A1), F-actin plus ends were tethered to 64 membrane-bound nodes, as shown for ring assembly (20, 43). In (A2), the plus end of each F-actin was tethered to a random membrane bead. In (A3), tethering could occur on any actin bead along the filament, and in (A4) F-actin was not tethered to the membrane. The four configurations of myosin are illustrated in Fig. 3 (panels M1–M4). In (M1), unipolar myosins were tethered by their tails to 64 membrane-bound nodes, again, as shown for ring assembly (20, 43). In (M2), unipolar myosins were tethered to the membrane in pairs. In (M3), each unipolar myosin was tethered to a random membrane bead and in (M4), myosins were modeled as bipolar molecules, randomly distributed throughout the ring, unattached to the membrane. The basic principles of constriction that were discovered are presented below.

**Figure 3:**
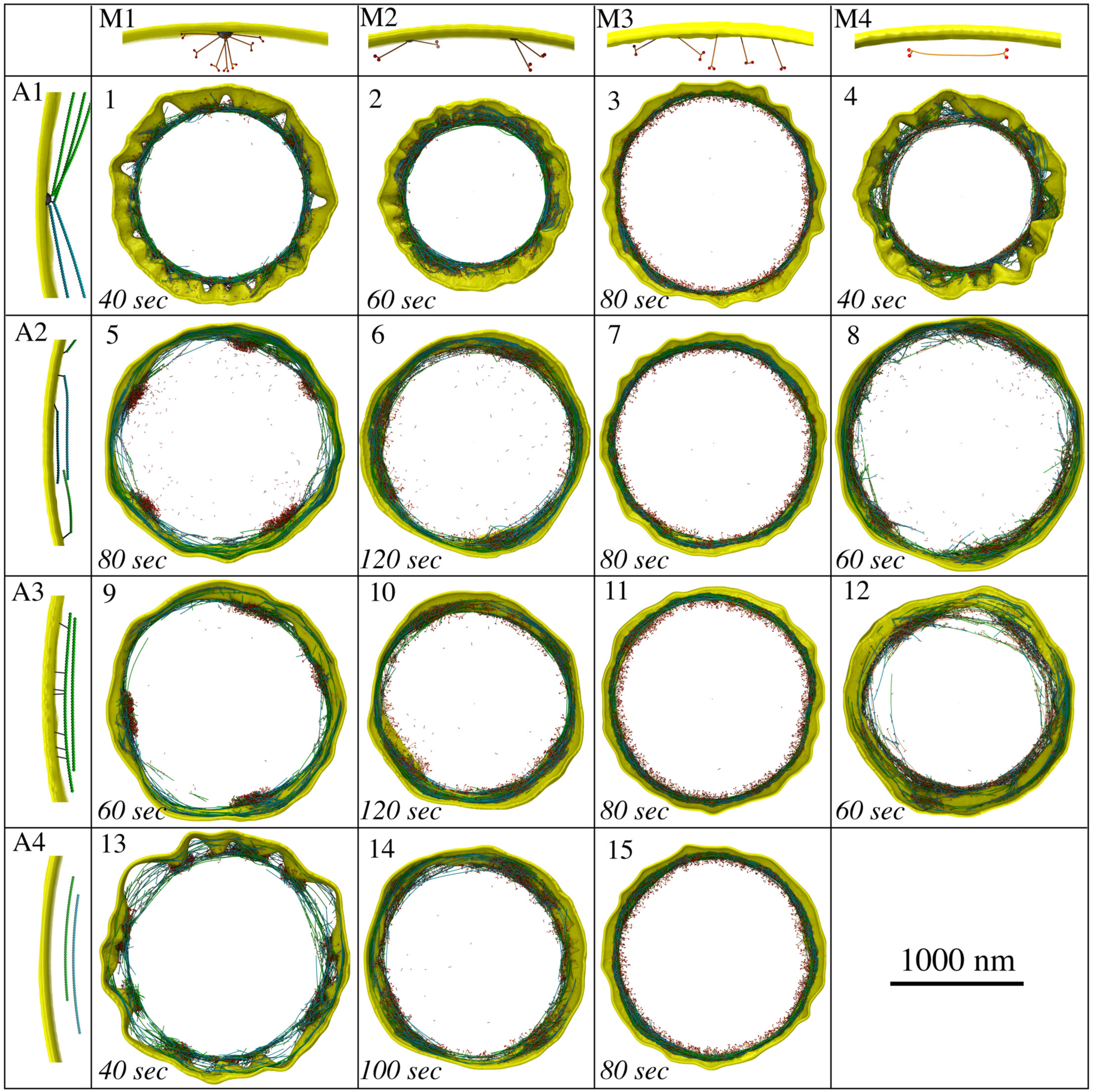
Eploration of different actomyosin models resulting from combining four actin configurations (A1–A4) with four myosin configurations (M1–M4). Resultant snapshots of the fifteen plausible models are presented. Note that the combination of A4 and M4 is not plausible since there are no tethers between the contractile ring and the membrane.

### Ring tension

First we calculated the ring tension of all the models (Fig. 4). In models where actin and myosins were anchored to pull on one another in a tug-of-war fashion (e.g., model 1–4 where actin was connected in nodes, model 3 being an exception), the ring produced a large tension. Meanwhile the ring produced a small tension if myosins were unipolar and individually attached to the fluidic membrane (models 3, 7, 11, 15). All models, however, produced tensions of similar order to the ring tension observed experimentally (24). This suggests that, at least within our models, comparison of the ring tension is not a definitive criterion to rule out certain models.

**Figure 4:**
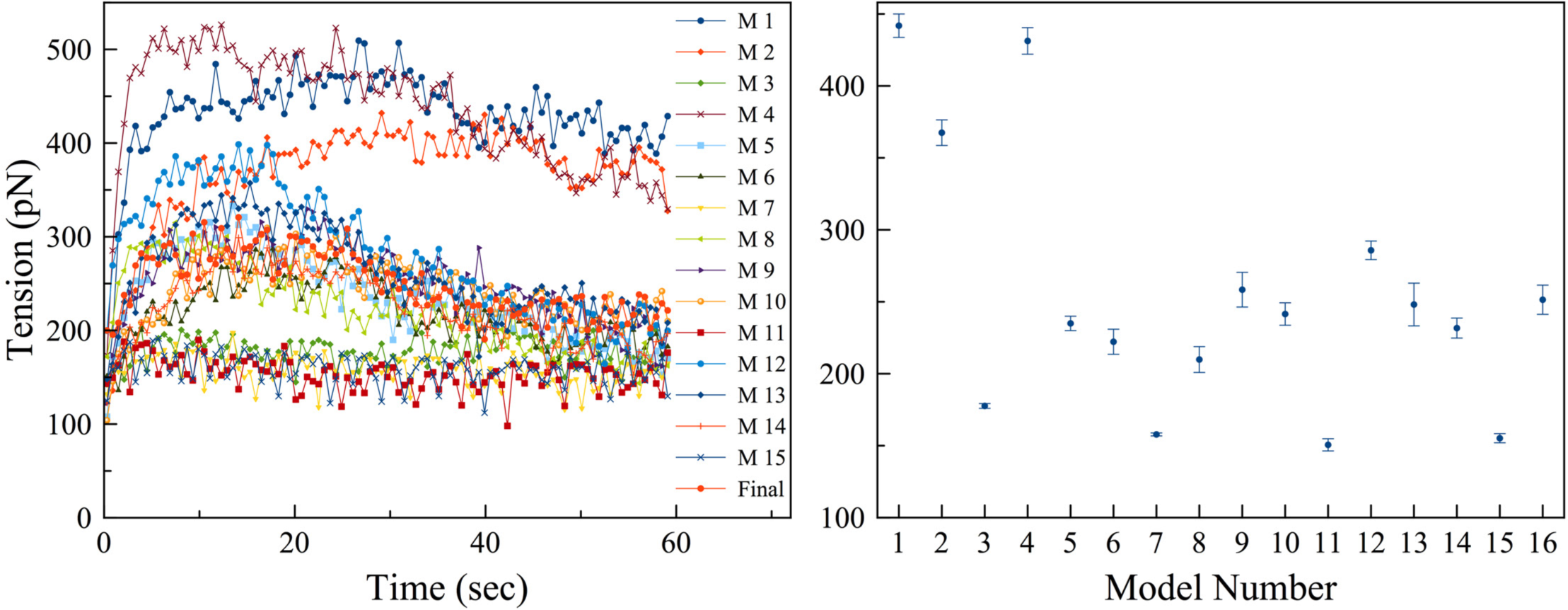
The ring tension was calculated: (left) representative individual time courses of the ring tensions, and (right) their averages over 5 simulations produced by 16 models with number 16 representing the final model. Error bars represent standard deviations.

### Individually, homogeneously distributed unipolar myosins maintain membrane smoothness

Several scenarios led to loss of membrane smoothness and circularity. One obvious cause was focusing the constriction force on only a small number of membrane sites. The most severe distortion occurred when the ring was connected to the membrane via only 64 nodes, as in models 1, 4, and 13, which resulted in membrane puckering during constriction (Fig. 3; SI Appendix/Fig. S2 & S3; Movie S1, at 7:08). As new cell wall material filled the gap between the membrane and the cell wall, puckering also occurred on the leading edge of the septum (SI Appendix/Fig. S2B, right panel), supporting the membrane puckers against turgor pressure. We found that our fluidic membrane model allowed nodes to slide (Fig. S4) with speeds comparable to those during ring assembly reported experimentally and via simulations (15, 37). As a result of node sliding, in several cases, puckers coalesced making large membrane deformations (Fig. S2C; Movie S1, at 7:50). Neither reducing the concentrations of actin, myosin, and crosslinkers in half (SI Appendix/Fig. S2C) nor doubling them (SI Appendix/Fig. S2D) mitigated puckering. The defects persisted even as the number of nodes increased from 64 to 140 (SI Appendix/Fig. S2E); the latter was recently reported by Laplante et al (27). We then studied how puckering depended on the mechanosensitivity of cell wall growth by varying *F_m_*, the minimal radial force on a membrane bead that induces cell wall growth (defined in **Methods/Cell wall and turgor pressure**). Increasing *F_m_* 100 times suppressed cell wall growth when unipolar myosins were individually connected to the membrane, but this low mechanosensitivity did not prevent nodes-induced puckering (SI Appendix/Fig. S5). Because the membrane in every cryotomogram appeared smooth (16), we know small puckers do not form in vivo, noting however that puckers larger than the 200 nm-thick cryosections cannot be ruled out. In our simulations, the presence of membrane puckers often caused actin filaments to lie at large angles with respect to the membrane (Fig. 5A; SI Appendix/Fig. S2B; S3B; S3C). By contrast, in other models which did not produce membrane puckers, filaments remained parallel to the membrane (Fig. 6), which is consistent with experimental observation (16). Smaller membrane puckers were observed in model 2, where unipolar myosins were attached to the membrane in pairs (Fig. 3; SI Appendix/Fig. S3A; Movie S1, at 8:11). On the other hand, in models 3, 7, 11 and 15, where unipolar myosin was individually attached to the membrane, providing an abundance of attachments, the membrane constricted without losing smoothness and actin filaments stayed parallel to the membrane (Fig. 3; Fig. 6; SI Appendix/Fig. S6; Movie S1, at 11:27). Therefore, if unipolar myosins exist during constriction, they are likely attached to the membrane individually.

**Figure 5:**
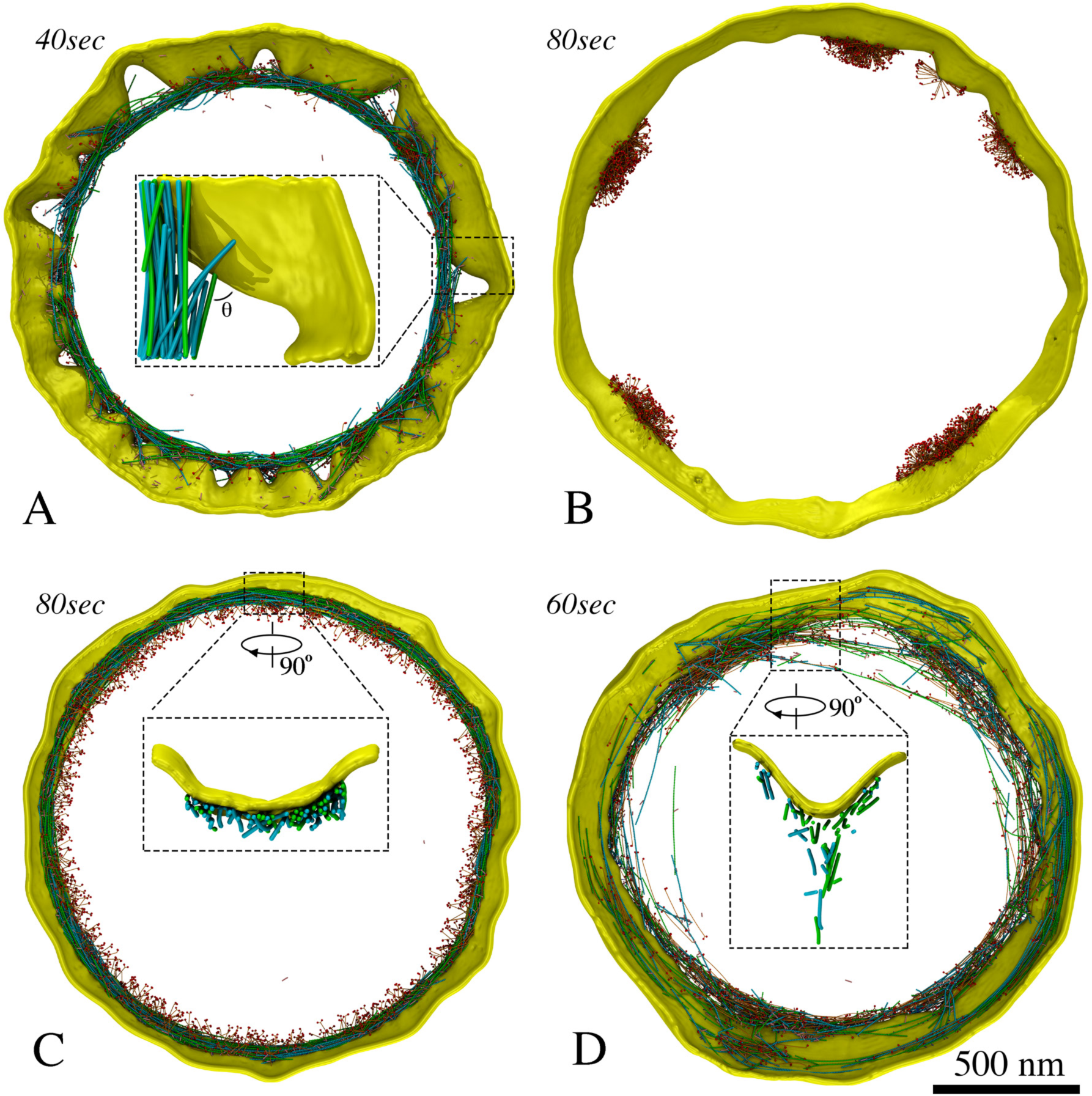
Representative features produced by the fifteen models. (A) Membrane puckering, as occurred in model 1, and large angles between filaments and membrane. (B) Tethering membrane-bound unipolar myosins in nodes or pairs, as in columns 1 and 2 of Fig. 3, resulted in aggregation. (C) Individual unipolar myosins, as in model 11, pulled filaments close to the membrane. (D) Bipolar myosins, as in model 12, pulled filaments away from the membrane.

**Figure 6:**
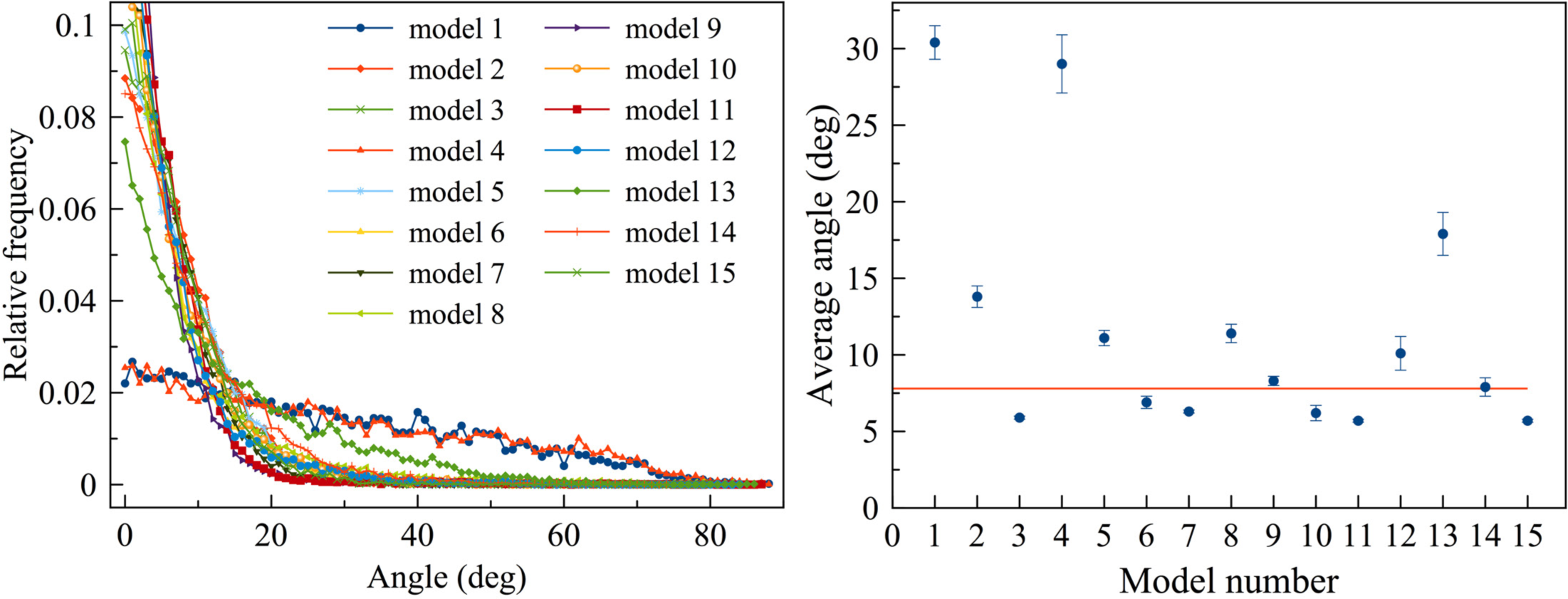
Angles between actin filaments and the membrane calculated after 60 sec of simulated time: (left) representative histograms of angles in individual simulations and (right) averages over five simulations for each model with error bars representing standard deviation and the red horizontal line indicating the average angle (7.8°) measured from electron tomograms for a reference. The presence of puckers, as in models 1, 4, and 13, causes filaments to form angles larger than observed experimentally.

Since a previous study observed that during ring assembly, actin and myosins in a broad band of nodes could coalesce into different structures when the crosslinker concentration varied (37), we explored whether changing the crosslinker concentration influenced the ring architecture in our simulations. Doubling or halving the crosslinker concentration did not change the ring architecture or basic outcome of any of our constriction models.

### Attaching unipolar myosin individually to the membrane prevents aggregation

Among models with abundant membrane attachments, in 5, 6, 9, 10 and 14 membrane deformation still occurred due to myosin aggregation. In contrast to fluorescence microscopy observations (44, 45, 22), myosins in these models gradually clumped together into a few large aggregates along the ring (Fig. 5B). Aggregation of unipolar myosins occurred through entanglement as either membrane nodes (models 5 and 9; Fig. 3; SI Appendix/Fig. S7; Movie S1, at 8:31) or pairs of myosins (models 6, 10 and 14; Fig. 3; SI Appendix/Fig. S8; Movie S1, at 9:12) became caught on each other due to steric hindrance while sliding along the membrane. Entangled myosin clusters were in turn larger, increasing the chance for further entanglement and creating a positive feedback that exaggerated the defect as constriction proceeded. As aggregation eventually concentrated the constrictive force, membrane circularity was lost. Varying the myosin turnover rate in models 6, 10, and 14, we found that myosin aggregation was mitigated when the myosin turnover rate was increased to 15 times faster or more than the rate we observed experimentally (SI Appendix/Fig. S9; Table S1). In model 8, where actin plus-ends were tethered to the membrane and bipolar myosin was not, clustering of plus-end tethers also led to myosin aggregation at these locations (Fig. 3; SI Appendix/Fig. S10; Movie S1, at 10:23). In contrast, in models 3, 7, 11 and 15, the uniform distribution of myosin provided a persistent, homogenous distribution of constrictive force that preserved membrane smoothness and circularity (Fig. 3; SI Appendix/Fig. S6; Movie S1, at 11:27) further supporting the notion that unipolar myosins are individually tethered to the membrane.

### Bipolar myosins pull actin filaments away from the membrane

Next, we focused on the five models where the membrane remained smooth (models 3, 7, 11, 12 & 15) and measured the distance between F-actin and the membrane (SI Appendix/Fig. S11). The four models containing individually tethered unipolar myosins (models 3, 7, 11 & 15) restricted filaments to ~21 nm from the membrane (Fig. 5C; SI Appendix/Fig. S6; Fig. S11), while ECT showed an average distance of ~60 nm (16). Due to membrane-tethering and pulling forces from the unipolar myosins, less than 0.2% of the actin beads in these four models were at a distance larger than 60 nm. In model 12, untethered bipolar myosins tended to pull actin away from the membrane, producing a larger average distance of 32 nm with nearly 10% of the actin beads at a distance larger than 60 nm (Fig. 5D; SI Appendix/Fig. S11). This suggested the presence of bipolar myosin within the ring in real cells. In some cases, actomyosin bundles consisting of unattached F-actin and bipolar myosins peeled off from the ring and depolymerized (Fig. 5D; Movie S1, at 10:51). This is consistent with previous observations by fluorescence microscopy (22), further supporting the presence of bipolar myosins and suggesting that actin filaments are not attached to the membrane.

### Final model: dual myosin configurations

We therefore built a final model consisting of untethered F-actin, individual unipolar myosins, and bipolar myosins (Fig. 7A). Simulating the final model resulted in normal constriction without visible defects of the membrane or ring (Fig. 7B; Movie S1, at 13:27). In this model, unipolar myosins remained at the outer edge of the ring due to their membrane attachment, while the bipolar form drifted inward, pulled toward the center by interaction with F-actin (Fig. 7C; Movie S2, at 13:54), matching the fluorescence microscopy result of the two myosin isoforms Myo2p and Myp2p (22). In our simulations, interactions with bipolar myosins pulled actin filaments away from the membrane (Fig. 7B, zoomed-in view; Fig. 7D), approximately recapitulating the distances observed by ECT (16), and occasionally caused actin/bipolar myosin bundles to peel off, as reported previously for actin/Myp2p bundles (22). Reducing the ATPase rate of the unipolar myosin in the simulation caused actin/bipolar myosin bundles to peel off more frequently, again in agreement with fluorescence microscopy results in which the loss of actin/Myp2p bundles occurred at higher frequency when the biochemical activity of Myo2p was reduced (22).

**Figure 7:**
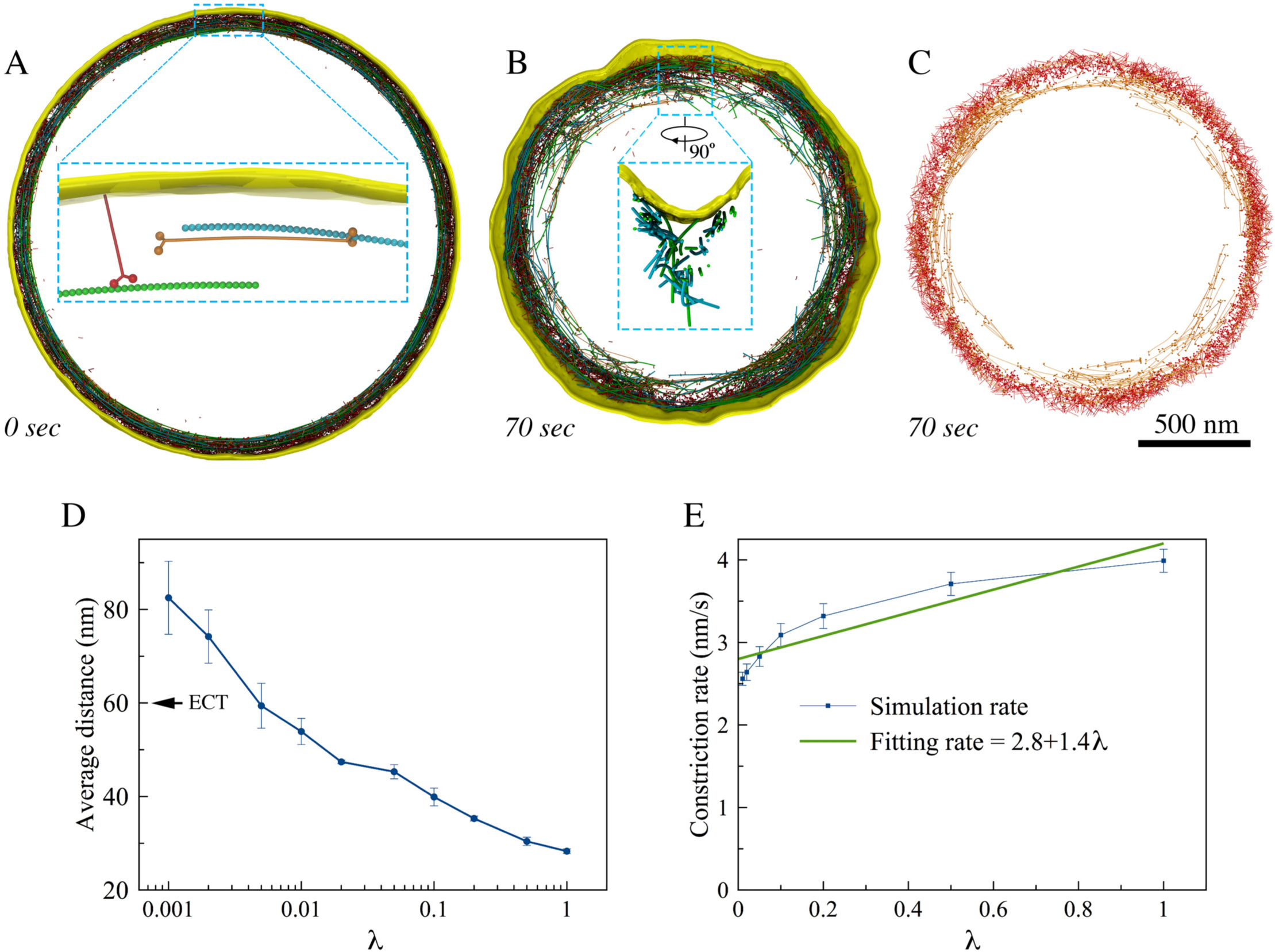
The final model. (A) A zoomed-in view shows initial configuration of the ring, including untethered F-actin (green and cyan), membrane-attached unipolar myosins (red), and bipolar myosins (orange). During constriction, (B) membrane smoothness and circularity were preserved and distances between F-actin and the membrane as observed in tomograms were recapitulated (as shown in a zoomed-in view), and (C) the membrane-attached unipolar myosins (red) occupied the outer edge of the ring while the unattached bipolar myosins (orange) occupied the inner edge. (D) Average distance between the simulated F-actin and the membrane (arrow indicates the average distance measured in tomograms) and (E) constriction rate as a function of the unipolar myosin's ATPase rate scaling factor, *λ*. Error bars represent standard deviations (n = 5).

Reasoning that the balance of force between unipolar myosins pulling F-actin close to the membrane and bipolar myosins pulling it away would dictate its average distance to the membrane, we investigated how the average distance between F-actin and the membrane depended on the ATPase rate of the unipolar myosin by scaling it with a factor *λ*. As expected, the average distance between F-actin and the membrane increased as the ATPase rate of the unipolar myosin decreased, reaching the experimentally measured value of 60 nm at *λ*~0.005 (Fig. 7D).

To further dissect the roles of the two forms of myosin, we studied the simulated constriction rate, *v* = Δ*r*/Δ*t*, defined as the ratio of average inward radial growth of the cell wall Δ*r* to constriction time Δ*t*, as a function of the unipolar myosin's ATPase rate (scaled with factor *λ*) (Fig. 7E). For simplicity, *v* was considered a linear combination of contributions from the bipolar myosin *v_b_* and the unipolar myosin *v_u_*. Fitting *v* = *v_b_* + *v_u_λ* to the simulated data yielded *v_b_* = 2.8 nm/s and *v_u_* = 1.4 nm/s. Since there were 2,000 bipolar and 3,200 unipolar myosin heads, on average, each bipolar head contributed an amount of ~1.4 pm/s to the constriction rate while each unipolar head contributed ~0.4 pm/s. The efficiency of the bipolar myosins in our simulations was therefore several times that of the unipolar myosins, likely due to the fact that unipolar myosins were attached to the fluidic membrane. This is in agreement with the experiments that showed Myp2p contributes more to the constriction rate of real cells than Myo2p (22).

## Discussion

From a methodological standpoint, we have demonstrated how 3D coarse-grained simulations can be used to explore complex models and hypotheses. The ring components were modeled in individual molecular detail, exerting force on a flexible membrane. Individual steps of myosin II's ATPase cycle were also modeled to produce power-stroke-driven movement of myosin along actin filaments. While we could not of course include all relevant molecules (only ~4 of the more than 100 proteins involved were modeled) or fully explore parameter space, our results did nevertheless suggest several interesting principles.

### The role of crosslinkers

Previous experimental studies have shown that crosslinkers such as α-actinin and fimbrin are essential for assembly of the fission yeast's AMR ring, but their role during constriction has not been clear (28–30). Earlier simulations showed that end-tracking crosslinkers and actin filament depolymerization could together drive contraction (36), but α-actinin and fimbrin are not end-tracking, and it remains unclear whether end-tracking crosslinkers are present in the ring. It was also previously suggested that contractility could arise in the presence of thick myosin filaments if they functioned as crosslinkers by remaining bound to the barbed end of F-actin (46). Although this might promote connectivity for long-range propagation of tension, it was unclear how such binding would be maintained, and thick myosin filaments were not seen in the cryotomograms (16). Our simulations suggest that crosslinkers like α-actinin and fimbrin allow long-range propagation of tension around the ring. This is consistent with findings on the contractility of *in vitro* ring-like (47) and disordered networks of actin (48).

### F-actin straightness

ECT revealed that F-actin filaments in dividing cells are remarkably straight (16). While in our first simulations involving only F-actin and myosin, the actin filaments became highly bent, here we identified two factors that likely reduce this bending *in vivo*. First, it has been shown *in vitro* that myosin binds preferentially to F-actins under tension (39). Biasing myosins to preferentially bind stretched F-actin filaments in our simulations reduced bending, and also helped maintain ring tension. It has also been shown *in vitro* that cofilin preferentially severs F-actins not under tension (40). Biasing cofilin's activity to bent filaments here promoted filament straightness. Our simulations therefore suggest one rationale for the otherwise puzzling presence in the ring of an actin severing factor (32, 33).

### Comparisons to previous simulations/treatments of actomyosin systems

Dasanayake et al. (49) studied 2D disordered networks of actin, myosin, and crosslinkers and found that they were by nature contractile, in agreement with our findings for the interplay of these three basic elements. Lenz also explored the behavior of disordered 2D networks, and found analytically that “contractile forces result mostly from motors plucking the filaments transversely” (50). The architecture of the AMR is very different, since the actins are parallel and bundled into a ring. As a result, contractile forces in our simulations arose from motors sliding parallel filaments past each other. Stachowiak et al. simulated a 2D actomyosin band where nodes containing 40 bipolar myosins each were modeled as single beads (24). The authors observed clustering of myosin beads when protein turnover was stopped, but the cause of aggregation was very different than seen here because in their model volume exclusion was not applied to all elements (e.g., objects could pass though actin filaments). In contrast, by modeling all the basic elements (including the membrane) in 3D and applying volume exclusion to all objects, we found aggregation occurred when actin filaments and unipolar myosins were connected to the membrane in nodes or pairs, throughout a range of physiologically relevant turnover rates. Further, our simulations allowed the characteristics and consequences of different actomyosin configurations to be assessed in 3D, and compared directly with those observed in cryotomograms (16). This revealed that concentrating force at nodes produces puckers in the membrane. Moreover, while Stachowiak et al.'s simulations produced tension similar to that measured in fission yeast protoplasts, our results showed that other actomyosin configurations can also produce ring tension of similar order. The most closely related previous work was that of Bidone et al., (37), who simulated how actin nodes placed on a 3D cylindrical surface can be drawn together into a tight ring by myosin filaments. The major difference with our work is that while Bidone et al. explored assembly of the ring, ours explored contraction, including changes in the shape of the cell wall boundary, and we found that concentrating force at nodes results in puckers. Thiyagarajan et al. simulated septum closure with a 2D representation of the cell wall (45). Assuming that the ring follows the shape of the septum leading edge, a condition we interpret as requiring an intimate and uniform connection to the membrane, and that the rate of cell wall growth was proportional to radial force, Thiyagarajan et al. showed that cell wall growth in local depressions would be faster than in flatter regions, and this could maintain circularity. This is most like our model in which myosins were connected to the membrane individually, since then force was distributed across thousands of connections, and in our case, this architecture also maintained circularity. Our simulation went on to show, however, that when force was concentrated at nodes, the basic assumption of uniform connection broke down and puckers resulted.

### Do nodes exist during constriction?

Actin filaments and myosins have been shown to form nodes during the assembly of the ring (11–15, 51), but a more recent study reported that the head domain of Myo2p distributed along pre-constriction rings (52), reflecting a discrepancy in the literature. While Laplante et al. recently suggested nodes persist during constriction (27), our results call into question whether this is the case. In our simulations, whenever constrictive force was concentrated on nodes or aggregates, membrane puckers formed, which is intuitively reasonable and we are not surprised this is invariant across crosslinker concentrations, myosin processivity, turnover rates, etc. As a consequence, large angles were frequently created between F-actin and the membrane (Fig. 5A, SI Appendix/Fig. S2B; S3B; S3C), features not seen in the cryotomograms (16). We conclude that either nodes are not present during constriction or we don't understand yet what other cellular forces maintain smooth membranes when constrictive force is concentrated at nodes. Perhaps future experiments will provide new insight into how membrane puckers are prevented.

### Actomyosin architecture

Instead of being directly attached to the membrane in nodes, our simulations suggest that actin filaments are not attached to the membrane. This rationalizes how bundles of actomyosin were able to separate from the membrane in fluorescence microscopy experiments (22). Our simulations also favored models where unipolar myosins link the ring and the membrane. While no clear evidence of such connections were seen in cryotomograms (16), the coiled-coil tail of a unipolar myosin is too thin and flexible to be resolved by ECT. Considering that Myo2p is the only myosin essential for viability (6, 21), it is a reasonable candidate for this role. Unipolar Myo2p molecules have already been proposed to attach to the membrane at nodes during ring assembly (20), but our results suggest they are more likely attached to the membrane individually to prevent aggregation and preserve membrane smoothness and circularity. Further, our results suggest that the myosin isoform Myp2p may exist in a bipolar configuration within the ring. This would explain fluorescence light microscopy experiments that showed that Myp2p primarily drives constriction, occupies the inner subdomain of the ring, and causes actomyosin bundles to peel away from the ring (22).

## Methods

For convenience, the key parameters of our simulations are listed in SI Appendix/Table S2.

### Actin filament

We modeled the actin filament (F-actin) as a chain of beads connected by springs (Fig. 1). Considering the double-helical nature of the filament, for convenience, each model bead represented two globular actin monomers (G-actin). Since 13 G-actins, corresponding to 6.5 model beads, cover a length of 35.9 nm (53), the relaxed length of the connecting spring is *l_a_* = 5.5 nm. The tensile modulus of F-actin has been measured to be *E* = 1.8 nN/nm^2^ (54). Estimating the cross-section of F-actin to be *A*~30 nm^2^ we derived the force constant of our model springs to be *k_a_* = *EA*/*l_a_*~10 nN/nm, reflecting that F-actin is not easily stretched. To reduce the computational cost of simulating such stiff springs, however, we used a force constant of 1 nN/nm considering the fact that the stretching of the F-actin was still negligible with this constant. To recapitulate actin's semi-flexibility, bending at a bead with an angle *θ* was penalized with an energy of 
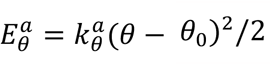
 where *θ_0_* = 180° was the relaxed angle, and the bending stiffness constant 
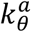
 was derived using the measured persistence length, *L_p_*~10 μm (55), to be 
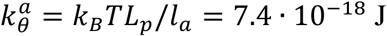
 where *k_B_* is the Boltzmann constant, and *T* = 295 K is the room temperature. Note that in initial simulations (see **F-actin straightness regulatory factors**) filaments became highly bent with the original bending stiffness 
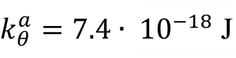
, but bending was prevented in the presence of straightness regulatory factors (SI Appendix/Fig. S1). Bending was also prevented even after 
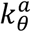
 was reduced three times to 2.4 ∙ 10^™18^ J, confirming that this reduction did not change the outcome of our simulations. Again, to reduce the computational cost, we then used 
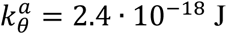
 for the rest of our simulations.

### Myosin configuration

Myosin was modeled to be either unipolar or bipolar and the same parameters were used for both configurations. Unipolar myosin was modeled as an 8-bead tail (representing the elongated C-terminal coiled-coil tail domain of two myosin heavy chains) connected to two head beads representing the N-terminal motor domains of the two heavy chains (Fig. 1). Bipolar myosin was composed of two unipolar molecules connected at the tails. Like the actin filament, the beads were connected by springs of force constant *k_m_* = 1 nN/nm, and relaxed length *l_m_* = 10 nm, which was chosen to reproduce a length of ~80 nm reported for the fission yeast conventional myosin II (56). To recapitulate the experimentally reported pulling force of 3–4 pN by a single myosin head (57), simulations were done where a unipolar myosin interacted with an actin filament from which the bending stiffness constant was determined to be 
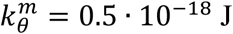
 (SI Appendix/Fig. S12). The relaxed angle was 180° on the tail, but at the head-to-tail junction it varied depending on the ATPase status of the head bead (see below for details).

### Myosin ATPase cycle

To model interaction with actin, each myosin head was allowed to exist in five phases: bound to (i) ATP, (ii) ADP and the hydrolyzed *P*_i_, (iii) ADP, *P*_i_ and actin, (iv) ADP and actin (*P*_i_ was released), and (v) actin (ADP was released). The relaxed angle at the head-tail junction was 120° if the myosin head was in phases (ii) or (iii) and 60° if in phases (i), (iv), or (v). Since ATPase rates for the individual phases of myosin II in fission yeast are not known, the probabilities of each phase transition were calculated based on studies from different species (58, 59). Specifically, ATP hydrolysis (phase (i) to (ii) transition) occurred with a probability of *p*_1_ = 25/s. If a myosin head in phase (ii) was within an interaction distance *D* = 15 nm from an unbound actin bead, actomyosin binding (phase (ii) to (iii) transition) occurred with a probability of *p*_2_ = 50/s. If there were more than one actin bead within *D*, the probability of being chosen for actin bead *i* was calculated as

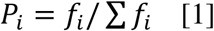

where 
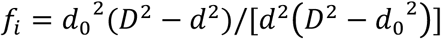
 was a function of the distance *d* between the myosin head and actin bead *i* and *d*_0_ = 5 nm was the relaxed distance between them once they were bound to each other. Myosin II is known to walk on F-actin directionally from the pointed end to the barbed end. To model this property, for simplicity, binding between myosin and actin was allowed only if the angle *θ* formed by the head-to-tail myosin vector and the plus-to-minus end actin vector was smaller than 90° (SI Appendix/Fig. S13A). Release of *P*_i_ (phase (iii) to (iv) transition) occurred with a probability of *p_3_* = 25/s, generating a pulling force in a power stroke fashion as the head-tail angle relaxed from 120° to 60°. ADP release (phase (iv) to (v) transition) occurred with a probability of *p_4_* = 25/s. Finally ATP binding and actin release (phase (v) to (i) transition) occurred with a probability of *p_5_* = 150/s. Our implemented rates of the myosin ATPase cycle resulted in an average myosin duty ratio of 
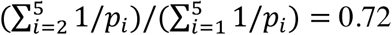
. While these rates set the upper limit of the load-free velocity of a myosin molecule to 
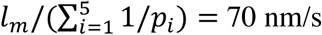
, a previous experimental study reported a myosin load-free velocity of 500 nm/s (60), reflecting a discrepancy in the literature.

### Actin Crosslinkers

Crosslinkers were modeled as two actin-binding domain (ABD) beads connected to a central bead by two springs of a force constant *k_c_* and relaxed length *l_c_* (Fig. 1). To account for the existence of different potential crosslinkers in real cells, namely *α*G-actinin and fimbrin (30), two types of crosslinkers were modeled. The one representing *α*-actinin had a length of 
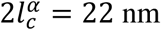
, the combined length of two ABDs (5 nm each) and two spectrin repeats (6 nm each) estimated from PDB structure 4D1E (while human *α*-actinin has four, *α*-actinin of fission yeast has only two spectrin repeats (61)), and 
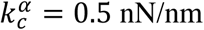
. The other representing fimbrin had 
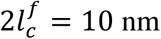
 (estimated from PDB structure 1RT8) and 
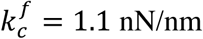
, which was chosen so that the two crosslinkers had the same Young's modulus, meaning 
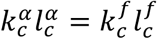
. To promote stiffness, bending with an angle *θ* was penalized with an energy of 
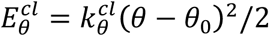
, where *θ*_0_ = 180° was the relaxed angle and the bending stiffness constant was 
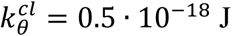
. Note that the spring constant for crosslinkers in our model was four orders of magnitude larger than that used in a previous simulation work by Stachowiak et al. (24) where the authors sourced an experimental work by Claessens et al. (62). In our opinion, Stachowiak et al. misinterpreted *k*_∥_ = 0.025 pN/nm (which was defined by Claessens et al. as the crosslinker's effective shear stiffness at very small deformations) as the crosslinker's extensional stiffness. Thermal forces would unrealistically stretch crosslinkers of this small spring constant tens of nm.

The binding of crosslinkers to actin was modeled to be stochastic. The binding of a crosslinker ABD bead to an actin bead within the interaction distance *D* = 15 nm occurred with a probability of 100/s. Similar to myosin-actin binding, if there were more than one actin bead within *D*, the probability of being chosen for actin bead *i* was calculated using equation [1]. Actin release from *α*-actinin and fimbrin occurred with probabilities of 3/s (63, 64) and 0.05/s respectively (28).

### Membrane

The membrane was modeled as a single layer of beads initially forming a cylinder (Fig. 1). To preserve membrane integrity, attractive forces were introduced between neighboring beads. To do this, a mesh of non-overlapping triangles with vertices on the beads was calculated from which non-redundant pairs of neighbor beads were determined. If a pair of beads were separated at a distance *d* larger than *D_pair_* = 20 nm, they were pulled together with a force of *F_pull_* = *k_pair_*(*d* − *d_pair_*)^2^ where *k_pair_* = 20 pN/nm^2^ was a force constant. To prevent the beads from being too close to each other, they were pushed apart with a force of *F_push_* = *k_pair_*(*d_mb_* − *d*)^2^ if *d* was smaller than a distance *d_mb_* = 10 nm. Since a permanent pairwise interaction would have prevented membrane beads from moving away from one another, blocking fluidity, the non-overlapping triangle mesh and therefore the non-redundant pair list were recalculated every 10^4^ steps. This allowed new pairs of beads to form based on their updated positions and made the membrane fluidic.

To generate membrane bending stiffness, a mesh of tetragons with vertices on the beads was calculated. If the four beads on each tetragon were not on the same plane such that the two diagonals were separated by a distance *d*, a spring-like force, *F_mb_* = *k_mb_d*, was exerted on the beads to pull the two diagonals towards each other (SI Appendix/Fig. S13B). Based on the reported membrane bending stiffness (65), the force constant was calculated to be *k_mb_* = 2 pN/nm. To prevent boundary artifacts, we applied a periodic boundary condition by translating the images of the beads of one edge to the other.

### Torque-facilitated crosslinker release

If two filaments were crosslinked at an angle *α* that was larger than 60° (SI Appendix/Fig. S13C) then once every 10^4^ time steps the crosslink was released with a probability *P_ux_* = 0.5 − *cos*(*α*).

### Cofilin function

If at an actin bead, the angle *α* between the tangent and the position vector from the barbed end (SI Appendix/Fig. S13D) was larger than 60°, once every 10^5^ time steps (the number was arbitrarily chosen since the rate in real cells is not known) the filament was broken into two segments with a probability *P_br_* = 1.0 − *cos*(*α*).

### Protein turnover

To model the turnover of ring components, actin depolymerization, addition of new F-actin, myosin removal and addition, and crosslinker removal and addition were included. At the beginning the G-actin pool was set empty for simplicity. Actin depolymerization was modeled to be stochastic, which removed an actin bead at the minus end to the G-actin pool with a probability of once every second, considering that F-actin turnover was reported to occur in about 1 min (41). A new filament of a randomly-selected length was added to a random location along the ring with a probability of once every 10^5^ time steps if the G-actin pool had more than 100 monomers. If membrane-bound nodes were present, the barbed end of the added F-actin was tethered to a random node.

A simple turnover mechanism was modeled for myosin. If all the heads of a myosin molecule were unbound, it was removed and a new one was added to a random location along the ring with a rate *r_t_* = 1/*τ*, where *τ* was the resident time of unbound myosins. For each model, we varied *τ* and measured the resultant average resident time of all myosins (bound and unbound). We report the resultant average resident times (SI Appendix/Table S1) that were close to 14 s, our experimentally-measured resident time (SI Appendix/Fig. S9), which is half of the previously reported value (41, 42). To explore the role of myosin turnover, multiple simulations of each model were run with different values of *τ*. The particular values used to produce each figure shown are listed in Table S1.

Similarly, to model crosslinker turnover, if both the ABD beads of a crosslinker were unbound, it was removed and a new one was added to a random location along the ring with a probability of once every 20 s (29).

### Protein binding force

If an actin bead and its binding partner (either a myosin head or a crosslinker ABD bead) were “bound” to each other at a given time step (see rules above for when they were considered bound), they exerted force on one another through a spring-like force *F_b_* = *k_b_*(*d* ™ *_0_*), where *k_b_* = 0.1 nN/nm was the force constant and *d*_0_ = 5 nm was the relaxed distance.

### Volume exclusion

To prevent the beads from overlapping with one another, if the distance *d* between any two beads was smaller than *r_off_* = 5 nm, they were pushed apart with a force *F_V_* = *K_V_*(*r_off_* −)^2^/(*d* − *r_on_*)^2^ to prevent them from approaching each other closer than *r_on_* = 4 nm, where *K_V_* = 0.1 nN.

### Membrane tethering

How tethering the ring to the membrane was modeled depended on the actomyosin configuration. In the node models, in which either F-actin plus ends or unipolar myosin tails (or both) were tethered to the membrane-bound nodes, each node was modeled as a bead connected to 10 nearest-neighbor membrane beads determined at the beginning. If the distance *d* between a node and a tethering counterpart, either an actin plus end, a unipolar myosin tail end, or a neighboring membrane bead, was larger than *d_n_* = 20 nm, the pair were pulled closer to each other with a force *F_n_* = *k_n_*(*d*− *d_n_*), where *k_n_* = 0.2 nN/nm was the force constant. In the paired-unipolar myosin configuration, for simplicity the two tail-end beads were tethered to a small node including 4 additional nearest-neighbor membrane beads. In the other models, direct tethering between one membrane bead to actin and/or unipolar myosin was modeled. If the distance *d* between an actin bead and its membrane tethering counterpart was larger than *d_t_* = 30 nm, the beads were pulled closer to each other with a force *F_t_* = *k_t_*(*d*− *d_t_*), where *k_t_* = 0.18 nN/nm was the force constant. If the distance *d* between a unipolar myosin tail-end bead and its membrane tethering counterpart was larger than *d_my_* = 5 nm, the beads were pulled closer to each other with a force *F_my_* = *k_my_*(*d*− *d_my_*), where *k_my_* = 0.2 nN/nm was the force constant.

### Cell wall and turgor pressure

Cell wall growth is needed to support ingression of the membrane since the tension from the AMR is not sufficient to counter the effect of large turgor pressure (66). Experiments have shown, however, that septum assembly slows down four folds (44, 66) and becomes misshapen in the absence of the contractile ring (16), suggesting ring constriction guides septum assembly in the normal condition. For simplicity, the membrane was treated as squeezable and the wall was modeled as a semi-rigid layer that expanded inwards following the membrane (SI Appendix/Fig. S14). The net force from turgor pressure and the cell wall on the membrane was modeled to follow Hook's law: a membrane bead at a distance *d* from the wall surface was pushed by a force *F_w_* = ™*k_w_*(*d*− *d_0_*), where *k_w_* = 0.05 pN/nm was the force constant and *d_0_* = 20 nm was the relaxed distance between the membrane and the wall. Previously, Thiyagarajan et al. proposed a tension-sensitive cell wall growth model in which the cell wall grows in proportion to the radial force exerted by the ring on the membrane (45). Similarly, to model cell wall growth, once every 10^3^ time steps, if the difference between *d* and *d_0_* was more than 0.1 nm (corresponding to a radial force of *F_m_* = 0.005 pN), the wall moved inward 0.01 nm.

Note that because it is not presently known what force would be required to initiate cell wall growth, this minimal radial force required to initiate cell wall synthesis (0.005 pN) was simply chosen as a value 20x smaller than the typical force from the ring (~ 0.1 pN). To explore the role of this mechanosensitivity parameter, simulations were also run with much larger *F_m_* values. We found that at *F_m_* = 0.5 pN (increased 100 times), there was essentially no cell wall growth in the model where unipolar myosins were individually connected to the membrane (distributing the ring constriction force homogeneously), but in the model where nodes were present, cell wall growth did occur, but puckers still formed (Fig. S5). Therefore, puckers were consistently the result of force concentration at nodes, not an artifact of a high mechanosensitivity.

### Diffusion

To model thermal motion of the system we introduced random forces on the beads. Each Cartesian component was generated following a Gaussian distribution using the Box-Muller transformation (67). Each transformation converted two random numbers from a uniform 0 – 1 distribution, *u_1_* and *u_2_*, into two random numbers of a Gaussian distribution:

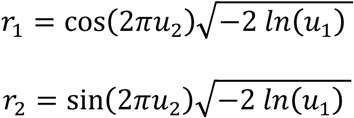

For a system of N particles, 3N/2 transformations were used to generate 3N numbers. While a pseudo random force can be generated by integrating a Gaussian random distribution with the time step, to reduce the computational cost, the random force was simply obtained by scaling the Gaussian random number with a force constant *k_r_*. To determine *k_r_* for actin we ran simulations of free individual actin filaments in the presence of the random force and compared the simulated tangent correlation, 〈*cosθ*〉, over distance *L* to the theoretical value *e^™L/*L_P_*^* where *L_P_* was the persistence length of the filament (SI Appendix/Fig. S15). We found that the simulated tangent correlation matched the theory best at *k_r_* = 20 pN. We then used the same *k_r_* = 20 pN for the random force on myosins and crosslinkers considering they were also cytoplasmic proteins. In the absence of relevant experimental measurements, we arbitrarily chose *k_r_* = 5 pN for the random force on the membrane.

### Initial ring configuration

To determine a minimal list of basic components of the ring, our model started with an actomyosin ring 200 nm wide (dimension along the long axis of the cell) and 30 nm thick (dimension along the radial direction) inside a membrane 300 nm wide and 1,000 nm in radius. The ring was composed of 400 F-actins of length chosen randomly in the range of 270 – 810 nm long (50 – 150 beads) resulting in ~30 – 40 filaments per ring cross-section, well within the range of 14 – 60 filaments observed by ECT (16)). 800 bipolar myosins were included. To study the role of crosslinkers, 600 *α*-actinins and 1,000 fimbrins were added to the ring. Note that these protein concentrations were within the ranges reported experimentally (68).

The same parameters for the membrane and crosslinkers were used for all 15 actomyosin configurations. The ring started 200 nm wide and 60 nm thick. Note that bundles of actomyosin peeled off the ring during constriction in model 12, where actin filaments were directly tethered to the membrane (see **Membrane tethering**) and myosin was bipolar, and this was also observed in the ring that started 30 nm thick. Either 800 bipolar (model 4, 8, 12) or 1,600 unipolar myosins (the other models) were present. The same ring configuration was used in simulations of the final working model except there were 1,600 unipolar and 500 bipolar myosins coexisting in the system. In all modeled rings, F-actin existed in two opposing polarities.

### Ring boundary

ECT showed that F-actins were strictly localized to the leading edge of the septum (16). This might be the result of either the ring tension or some physical barrier that was not distinguishable in the tomograms or both. The septin cytoskeletal proteins were thought to serve as such a barrier as they form a pair of rings flanking the actomyosin ring during constriction (69). This proposal was challenged later as the septin rings were reported to be dispensable for cytokinesis in budding yeast (70). In addition, the barrier function of septins is unlikely in fission yeast since the two rings do not contract during contraction of the actomyosin ring (12, 71, 72). Another barrier candidate, if required at all, could be the F-BAR protein Cdc15, as it was reported to form long filaments likely wrapping around the division site several times (73). This stable scaffold might restrict movement of partner proteins in the ring. To implement a diffusion barrier in our model, if a ring component bead moved a distance Δ*x* outside the ring boundary, chosen to be 200 nm wide along the ring axis, it was simply pulled back with a force of *k_br_*Δ*x*, where *k_br_* = 10 pN/nm was the force constant.

### System dynamics

To track the evolution of the system we used a simple molecular dynamics simulation. Specifically, the coordinate *X*(*t*) of each bead changed following the Langevin equation:

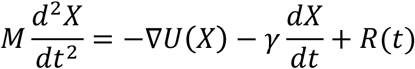

where *M* is the mass of the bead, *U* the interaction potential *γ* = 6 ∙ 10^™6^ Ns/m the damping constant and *R* the random force on the bead (see **Diffusion** above). To select a large damping constant that made simulations computationally efficient, we ran simulations where a single myosin molecule walked on a fixed actin filament and characterized the myosin load-free velocity with respect to the damping constant (SI Appendix/Fig. S16). A damping constant of *γ* = 6 ∙ 10^™6^ Ns/m was chosen to minimize computational cost without perturbing the myosin load-free velocity. Since we used the same damping constant for every bead in the system, the constant for a complex was proportional to the number of beads in the complex. Thus a small node of ~7 unipolar myosins (having ~70 beads) experienced a damping constant of ~420 pNs/μm, corresponding to a diffusion constant of ~10 nm^2^/s, the experimental value reported by Vavylonis et al (15). Assuming the inertia of the bead was negligible, and thus *M* = 0, the displacement was simply a linear function of total force *F*:

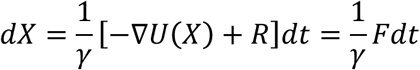

To prevent a large force from moving a bead too far, we constrained the maximal displacement of any bead in any time step (corresponding to the maximal force *F_max_*) to *D_max_* = 0.01 nm. Displacement *D* of each bead was then calculated as

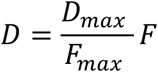

Since the time step was not a constant in our simulations, the average time step was calculated at the end of each simulation, which fell in the range of 0.2 – 0.3 μs. Simulation codes were written in Fortran and the trajectories of each system were visualized using VMD (Visual Molecular Dynamics) (74).

### F-actin straightness

To compare actin filament straightness in the tomograms and the simulations, we defined “straightness” as the filament's contour length *L_contour_* divided by the length of a straight line connecting the two ends *L_end–to–end_* (SI Appendix/Fig. S13E). Note that we did not compare persistence length, which is usually used to characterize free filaments not being pulled or acted upon by anything other than random thermal forces.

### Actin-membrane distance

To compare the distances between the actin filaments and membranes in the tomograms and the simulations, we defined the distance from an actin bead to the membrane as the smallest distance from the actin bead to any membrane bead.

### Constriction rate

For simplicity, the constriction rate was calculated as the inward growth of the cell wall, Δ*r*/Δ*t*, averaged around its circumference, where Δ*r* was the radial displacement of the cell wall leading edge and Δ*t* was the duration of constriction.

### Ring tension

To calculate the ring tension during constriction, first the ring radius *R_r_* was calculated as the average distance from the actin beads to the cell axis. The ring tension was then calculated as

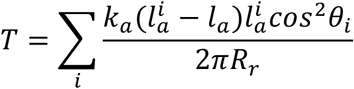

where the sum was over all actin springs *i* which had their length 
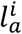
 larger than the relaxed length *l_a_*,*k_a_* was the actin spring constant, and *θ_i_* was the angle spring *i* deviated from the circumferential direction.

## Experimental procedures

### Microscopy

Mid-log-phase cells were spotted on a 2% Agar pad supplemented with YES media and observed under a custom-built spinning disk confocal microscope with an inverted Olympus IX-83,100X/1.4 plan-apo objective, a deep cooled Hamamatsu ORCA II ‐ER CCD camera and Yokogawa CSU:X1 spinning disk (Perkin-Elmer). A stack of 18–20 Z slices of 0.3 mm Z-step-size was collected every 2 min for an hour at 25°C using the Velocity software (Perkin-Elmer). Images were then rotated and cropped using the imageJ software to align cells and 3D reconstruction was done using the Velocity software.

### Fluorescence Recovery After Photobleaching (FRAP)

Cells were mounted onto a 2% Agar pad supplemented with YES media and observed under a Leica TCS SP8 scanning confocal microscope with a 63x magnification, 1.4 numerical aperture (NA) oil-immersion objective. The experiments were performed at 25°C unless otherwise indicated. For excitation of GFP, we used a 488 nm Argon laser. Images were collected with a scan speed of 40 fps, 12x digital zoom, at 256 x 256 pixels. The laser intensity for photobleaching was adjusted to obtain ~80% loss of fluorescence in the approximately 0.2 μm x 0.2 μm circular bleached region of the cytokinetic ring. To allow rapid bleaching, we used a high laser intensity with 1–3 iterations of the bleaching scan. The images were collected before and after bleaching, using low laser intensities and FRAP was monitored for 1.5 to 2 min. Data from the experiment were analyzed using ImageJ (National Institute Of Health, Bethesda, MD) with FRAP plugin www.embl.de/eamnet/frap/FRAP6.html) using the double normalization method (75). Normalized curves were fitted to single exponential functions to extract the mobile fraction and half-life.

## Acknowledgments

The authors thank Catherine Oikonomou for helping revise the manuscript for clarity. The work was supported in part by NIH grant R35 GM122588 to GJJ.

